# Benchmarking taxonomic classifiers with Illumina and Nanopore sequence data for clinical metagenomic diagnostic applications

**DOI:** 10.1101/2022.01.11.475979

**Authors:** Kumeren N. Govender, David W. Eyre

## Abstract

Culture-independent metagenomic detection of microbial species has the potential to provide rapid and precise real-time diagnostic results. However, it is potentially limited by sequencing and classification errors. We use simulated and real-world data to benchmark rates of species misclassification using 100 reference genomes for each of ten common bloodstream pathogens and six frequent blood culture contaminants (n=1600). Simulating both with and without sequencing error for both the Illumina and Oxford Nanopore platforms, we evaluated commonly used classification tools including Kraken2, Bracken, and Centrifuge, utilising mini (8GB) and standard (30-50GB) databases. Bracken with the standard database performed best, the median percentage of reads across both sequencing platforms identified correctly to the species level was 98.46% (IQR 93.0:99.3) [range 57.1:100]. For Kraken2 with a mini database, a commonly used combination, median species-level identification was 79.3% (IQR 39.1:88.8) [range 11.2:100]. Classification performance varied by species, with E. coli being more challenging to classify correctly (59.4% to 96.4% reads with correct species, varying by tool used). By filtering out shorter Nanopore reads (<3500bp) we found performance similar or superior to Illumina sequencing, despite higher sequencing error rates. Misclassification was more common when the misclassified species had a higher average nucleotide identity to the true species. Our findings highlight taxonomic misclassification of sequencing data occurs and varies by sequencing and analysis workflow. This “bioinformatic contamination” should be accounted for in metagenomic pipelines to ensure accurate results that can support clinical decision making.

**Importance:** Metagenomics may transform clinical microbiology by enabling more rapid species detection in a potentially unbiased manner and reducing reliance on culture-based approaches. However, it is still limited by ongoing challenges such as sequencing and classification software errors. In this study, we use simulated and real-world data to define the intrinsic rates of species misclassification that occur using Illumina and Oxford Nanopore sequencing platforms with commonly used taxonomic classification tools and databases. We quantify the extent of “bioinformatic contamination” arising from the classification process. This enables us to identify the best performing tools that maximize classification accuracy, and to suggest how taxonomic misclassification can be formally accounted for in clinical diagnostic workflows. Specifically, we specify thresholds for identifying or excluding polymicrobial infections in metagenomic samples, based on rates of misclassification of similar species, which might have clinical implications when treating infection.

## Background

Reliable identification of bacterial species is a fundamental part of clinical microbiology. Traditionally identification has been based on growth on selective media, morphological appearances, and biochemical tests. However, many microbiology laboratories now rely largely on MALDI-TOF mass spectrometry of cultured isolates to identify most species. In clinical metagenomics, a culture-independent approach is used to identify species present in a sample based on unselective DNA sequencing.

Multiple studies have compared the potential of metagenomic approaches to replicate traditional workflows, with sensitivity for species detection, regarding culture-based approaches as the reference standard, varying from 63% to 98% and specificity from 35-100% (1). Metagenomic species identification depends on the quality and properties of the sequence data used, e.g., long-read vs. short-read, the taxonomic classification algorithm and the database used to match sequences to species. Additionally approaches need to account for contamination while maximizing sensitivity and specificity. Contamination with human DNA is expected and can also arise from imperfect sampling, the laboratory environment, equipment, and reagents, and also bioinformatically. Bioinformatic contamination can occur from errors in demultiplexing where multiple samples are run at once using different indexes, and from misclassification of sequence data as the wrong genus or species.

One of the primary challenges in metagenomics is reads may be generated from multiple species rather than just one following sequencing of an isolate from culture. The creates a computational challenge, i.e., how to accurately identify and differentiate all species contained within a sample? This must also be done against an ever-increasing number of previous sequences to compare with. Metagenomic classifiers can be grouped based on their reference database type which include: DNA-to-DNA methods for example, Kraken (2), Kraken2 (3), Bracken (4) and PathSeq (5); DNA-to-protein methods for example, Kaiju (6) and Diamond (7); and DNA-to-marker methods which include only specific gene families in their reference databases for example, MetaPhlAn (8) and mOTUs (9). In a recent benchmarking study,(10) DNA-to-DNA methods were the best-scoring methods, while DNA-to-marker methods performed more poorly.

Some studies have benchmarked classifiers and observed that the performance of different classifiers varies significantly even on the same benchmark datasets (10–15). However, performance depends both on the taxonomic classifier and the reference database of sequences of known species used and few studies have compared performance using a uniform database(10). Similarly, there are also relative few studies using long-read sequencing data (16). Nevertheless, there is a need for thorough benchmarking to shed light on the optimal selection of bioinformatic tools for metagenomics and to account for “bioinformatic contamination” which is an important and potentially underrecognized issue in the field (17).

Here we use simulated and real-world data to define the intrinsic rates of species misclassification that occur using Illumina and Oxford Nanopore sequencing platforms with commonly used classification tools and databases. This enables us to quantify the extent of bioinformatic contamination arising from the classification process to guide the selection of optimal tools that maximize classification accuracy, and to formally account for bioinformatic contamination in clinical diagnostic workflows. Additionally, this allows us to specify thresholds for identifying or excluding polymicrobial infections in metagenomic samples.

## Methods

### Simulated data

We generated simulated sequence data for ten common blood stream pathogens (*Staphylococcus aureus, Streptococcus pneumoniae, Streptococcus pyogenes, Enterococcus faecalis, Pseudomonas aeruginosa, Escherichia coli, Klebsiella pneumoniae, Proteus mirabilis, Bacteroides fragilis* and *Enterobacter cloacae*) and six common blood culture contaminants (*Staphylococcus epidermidis, Staphylococcus haemolyticus, Bacillus cereus, Cutibacterium [Propionibacterium] acnes, Streptococcus mitis* and *Micrococcus luteus*). For each species, 100 randomly-selected reference genomes were downloaded (on the 10^th^ of January 2021) from the NCBI RefSeq database (18), i.e. 1600 genomes in total.

Simulated Illumina and Oxford Nanopore reads were generated from each genome, with separate simulations with and without sequencing errors. Illumina reads were simulated using ART_Illumina v2.3.7 (19) (art_illumina -i reference.fna -p -l 250 -f 800 -m 999 -s 1 -o -nf 5 - na -ss MS -na) generating 12 million paired fastq reads for each reference genome. Oxford Nanopore reads were simulated using NanoSim-H v1.1.0.4 (20, 21) (nanosim-h reference.fna -p ecoli_R9_1D -o -n 20000), generating 20,000 reads per genome. The default ecoli_R9_1D error profile and MiSeq error profile was used for Nanopore and Illumina read simulation respectively. No pre-processing of fastq reads were performed.

### Taxonomic classifiers and sequence databases

Simulated reads were used to benchmark three commonly used taxonomic classifiers: Kraken2 (v2.0.7) (3), Bracken (v2.5) (4) and Centrifuge (1.0.4) (22). Mini (8GB) and standard (30-50GB) uniform databases were built from archaeal, bacterial, viral and human reference genomes downloaded from NCBI RefSeq on the 10^th^ of January 2021. Databases were built according to each tool’s default setting. The mini database resembled an up-to-date version of “MiniKraken” limited to 8GB, which down-samples the standard Kraken2 database using a hash function. As Centrifuge does not allow for the restriction of size when building a database, only a standard database was built. Fastq files were classified using the default settings of Kraken2 (kraken2 --db $DBNAME sequence_data.fq --report /), Bracken (bracken -d $DBNAME -i sequence_data.fq -o /) and Centrifuge (centrifuge -k 1 -x $DBNAME -f / --report-file / -S /) however, for Centrifuge we set the distinct primary assignment value to 1 (-k 1) in order to provide a comparable standard where each read is assigned to only one taxonomic level. All classifier results were converted into a Kraken-style report for comparison.

### Real-world data

We also used real-world Illumina sequencing data for benchmarking, which consisted of reads generated from sequencing of 100 pure isolates of *Staphylococcus aureus, Escherichia coli* and *Klebsiella pneumoniae* from previous published works [NCBI project accession numbers PRJNA690682, PRJNA604975] (23, 24). Sequenced isolates originated from blood stream infections between 15-September-2008 and 01-December-2018 in Oxfordshire, UK. Available 150 bp paired-end short-read sequences were generated using the Illumina HiSeq 4000.

### Statistical analysis

We measured classifier performance as the proportion of reads classified as the correct species or correct genus, given the known species of the reference genome or clinical isolate from which reads were simulated or obtained. The reported read abundance for each species may be referred to as either “raw” calculated from the relative abundance of reads from each taxa or “corrected” by accounting for genome size, such that the relative number of organisms of each taxa is estimated. We used raw abundance estimates unless the correction is automatically performed by the software as in the case of Bracken.

We used generalized linear regression (with a logit link function, binomial family and robust standard errors) to investigate associations between the proportion of reads classified as the correct species and sequencing technology, taxonomic classifier, database and species. We also use these models to present probabilities of reads being classified correctly for specific combinations of these covariates.

Similarly, to investigate the extent to which species misclassification was because the true and misclassified genomes were similar, we used a generalized linear model to investigate the relationship between the average nucleotide identity (ANI) between the true species and the misclassified species and the proportion of reads misclassified as the incorrect species. ANI values for each correct-incorrect species pair were generated by taking the median value for all pairwise comparisons between 30 genomes of each species misclassified (downloaded from NCBI RefSeq on the 24^th^ June 2021) and 50 genomes of the correct species using FastANI (fastANI --ql [QUERY_LIST] --rl [REFERENCE_LIST] -o [OUTPUT_FILE]) (25). Analyses were performed using R (v.4.1) and R Studio (v1.4.1717) (26).

To investigate the relationship between read lengths and species classification performance, we generated Oxford Nanopore and Illumina reads of varying lengths using error and error-free profiles from 10 reference genomes of *Staphylococcus aureus* and *Escherichia coli* each, and plotted read lengths vs. the proportion of reads correctly classified by Kraken2.

## Results

### Determinants of taxonomic classification accuracy

We simulated sequence data from 1,600 genomes representing 10 common pathogens and 6 common blood culture contaminants. We chose *Staphylococcus aureus*, classified with Kraken2, using a mini (8gb) database, with simulated Illumina reads including sequencing errors as our reference category for comparisons as this is a common pathogen and common metagenomic workflow (Tables 1, and S1). Using this combination, 89.6% (95% CI, 88.4%:90.7%) of reads were classified as the correct species, and 96.9% (95.3%:97.7%) as the correct genus. We also present performance in probabilities in Table 2 and Table S2.

**Table 1.** Logistic regression model for performance of species abundance classification by sequencer, classifier, database and bacterial species.

|  |  | **Descriptive: Median (IQR)** | **Odds ratio** | **95% CI** | **p-value** |
| --- | --- | --- | --- | --- | --- |
| **Multivariable reference** | Species abundance classification (Reference: *Staphylococcus aureus*, Illumina with error profile, mini database, Kraken) | 89.6 (88.4:90.7) | [Reference] | - | - |
| **Sequencing profile** | Illumina with error profile [Reference] | 92.4 (60.1:97.8) | [Reference] | - | - |
|  | Illumina with error-free profile | 92.9 (71.8:97.5) | 1.01 | 0.96, 1.07 | 0.7 |
|  | Nanopore with error profile | 92.1 (74.9:97.3) | 1.10 | 1.04, 1.16 | <0.001 |
|  | Nanopore with error-free profile | 97.9 (93.9:99.3) | 2.99 | 2.80, 3.20 | <0.001 |
| **Database** | Mini database | 91.7 (73.2:98.1) | [Reference] | - | - |
|  | Standard database | 96.1 (83.9:98.5) | 2.02 | 1.92, 2.12 | <0.001 |
| **Classifier** | Kraken | 89.0 (70.3:96.8) | [Reference] | - | - |
|  | Bracken | 97.6 (90.9:99.0) | 3.06 | 2.92, 3.21 | <0.001 |
|  | Centrifuge | 94.8 (72.4:97.5) | 1.04 | 0.97, 1.10 | 0.3 |
| **Species** | *Cutibacterium acnes* | 99.6 (98.4:99.8) | 2.20 | 1.99, 2.43 | <0.001 |
|  | *Enterococcus faecalis* | 98.1 (96.2:99.1) | 1.41 | 1.33, 1.49 | <0.001 |
|  | *Staphylococcus epidermidis* | 97.4 (94.5:98.6) | 0.74 | 0.65, 0.84 | <0.001 |
|  | *Proteus mirabilis* | 97.6 (95.2:98.8) | 0.89 | 0.80, 1.00 | 0.058 |
|  | *Staphylococcus aureus* | 98.5 (93.3:99.3) | [Reference] | - | - |
|  | *Streptococcus pyogenes* | 97.7 (95.0:99.1) | 0.79 | 0.74, 0.84 | <0.001 |
|  | *Staphylococcus haemolyticus* | 96.7 (93.7:98.0) | 0.79 | 0.74, 0.85 | <0.001 |
|  | *Streptococcus pneumoniae* | 96.5 (93.8:98.5) | 0.59 | 0.55, 0.62 | <0.001 |
|  | *Bacteroides fragilis* | 95.3 (91.5:97.6) | 0.54 | 0.50, 0.58 | <0.001 |
|  | *Klebsiella pneumoniae* | 89.6 (70.0:93.7) | 0.13 | 0.13, 0.14 | <0.001 |
|  | *Escherichia coli* | 89.2 (73.3:95.8) | 0.17 | 0.16, 0.18 | <0.001 |
|  | *Micrococcus luteus* | 88.3 (59.8:96.0) | 0.15 | 0.14, 0.17 | <0.001 |
|  | *Pseudomonas aeruginosa* | 87.5 (75.1:95.8) | 0.16 | 0.15, 0.18 | <0.001 |
|  | *Streptococcus mitis* | 55.4 (37.9:72.3) | 0.04 | 0.04, 0.04 | <0.001 |
|  | *Bacillus cereus* | 34.1 (15.7:52.0) | 0.02 | 0.02, 0.02 | <0.001 |
|  | *Enterobacter cloacae* | 10.1 (3.0:92.6) | 0.02 | 0.02, 0.02 | <0.001 |

**Table 2.** Predicted abundance classification for *Staphylococcus aureus* and *Escherichia coli*.

| Sequencer | Error profile | Database | Classifier | Predicted performance (95% CI) |
| --- | --- | --- | --- | --- |
| *Staphylococcus aureus* |  |  |  |  |
| Nanopore | Error-free | Standard | Bracken | 99.4 (99.3:99.5) |
| Nanopore | Error | Standard | Bracken | 98.4 (98.2:98.5) |
| Illumina | Error | Standard | Bracken | 98.2 (98:98.4) |
| Nanopore | Error | Mini | Bracken | 96.7 (96.4:97.1) |
| Illumina | Error-free | Mini | Bracken | 96.5 (96.1:96.8) |
| Nanopore | Error-free | Mini | Centrifuge | 96.5 (96:96.9) |
| Nanopore | Error-free | Mini | Kraken | 96.4 (95.9:96.7) |
| Nanopore | Error | Standard | Kraken | 95.1 (94.6:95.6) |
| Illumina | Error | Standard | Kraken | 94.7 (94.1:95.2) |
| Nanopore | Error | Mini | Kraken | 90.7 (89.7:91.6) |
| Illumina | Error | Mini | Centrifuge | 90.1 (89:91.2) |
| Illumina | Error-free | Mini | Kraken | 89.9 (88.9:90.9) |
| Illumina | Error | Mini | Kraken | 89.8 (88.7:90.8) |
| *Escherichia coli* |  |  |  |  |
| Nanopore | Error-free | Standard | Bracken | 96.4 (96.2:96.7) |
| Nanopore | Error | Standard | Bracken | 90.8 (90.2:91.4) |
| Illumina | Error | Standard | Bracken | 90 (89.3:90.7) |
| Nanopore | Error | Mini | Bracken | 83.1 (82.1:84) |
| Illumina | Error-free | Mini | Bracken | 81.9 (80.8:82.9) |
| Nanopore | Error-free | Mini | Centrifuge | 81.9 (80.6:83.1) |
| Nanopore | Error-free | Mini | Kraken | 81.4 (80.3:82.4) |
| Nanopore | Error | Standard | Kraken | 76.4 (75.2:77.6) |
| Illumina | Error | Standard | Kraken | 74.7 (73.3:76) |
| Nanopore | Error | Mini | Kraken | 61.6 (60.1:63.2) |
| Illumina | Error | Mini | Centrifuge | 60.2 (58.3:62.1) |
| Illumina | Error-free | Mini | Kraken | 59.7 (58:61.3) |
| Illumina | Error | Mini | Kraken | 59.4 (57.7:61) |

In a multivariable model, using Oxford Nanopore (including with sequencing errors) independently improved the odds of classifying reads correctly to the species, while reducing the odds of classifying reads to the genus level (odds ratio vs. Illumina sequencing 1.10 (95% CI, 1.04:1.16) and 0.76 (0.72:0.80) respectively). As expected, simulating data without sequencing errors improved performance for both Illumina and Oxford Nanopore, however this level of performance cannot be currently achieved in reality (Table 1).

Use of the “standard”, i.e., larger, database independently improved species and genus level classification (odds ratio vs. Mini-database 2.02 (1.92:2.12) and 3.17 (3.03:3.32) respectively). Compared to Kraken2, Centrifuge performed similarly for species and inferiorly for genus classification (odds ratio vs. Kraken2 1.04 (0.97:1.10) and 0.70 (0.66:0.74) respectively), while Bracken performed superiorly for both species and genus (odds ratio vs Kraken 2 3.06 (2.92:3.21) and 6.29 (5.95:6.65) respectively).

Classification performance varied by species. Some species, including *Cutibacterium acnes* and *Enterococcus faecalis* had a higher adjusted probability of reads being correctly classified at the species level, compared to *S. aureus*, ranging between 92.5% to 99.7% across different sequencers, classifiers and databases (Table S2) (odds ratios ranging between 1.41 and 2.20 (Table 1)). Whereas other species were more challenging to classify correctly, e.g. *E. coli* with a predicted adjusted probability of reads being classified correctly ranging between 59.4% to 96.4% (odds ratio vs. S. aureus 0.17 (95%CI, 0.16:0.18)). Of note, *Streptococcus mitis, Bacillus cereus* and *Enterobacter cloacae* had seemingly poor performance ranging between 26.1%-86.8%, 14.1%-75.2% and 15.8%-77.6%, however, these species belong to a group or complex of bacteria which makes genomic differentiation more complex (27–29).

### Best performing combinations

Overall, Bracken with the standard database was the best performing option (Figure 1 and S1 show abundance classification details of five important common blood stream pathogens and five contaminants respectively). Across all 1,600 genomes, median performance by Bracken using a standard database for species identification was 98.46% (IQR 93.0:99.3) [range 57.1:100] and for genus was 99.6% (IQR 98.9:99.9) [range 34.4:100.0], and had no unclassified reads. Median performance, with a common combination, Kraken2 with a mini database, for species was 79.3% (IQR 39.1:88.8) [range 11.2:100] and for genus was 94.9% (IQR 87.6:98.0) [range 4.9:100.0], with a median of 0.9% (IQR 0.0:8.5) [range 0.0:14.3] unclassified reads.

**Figure 1.**
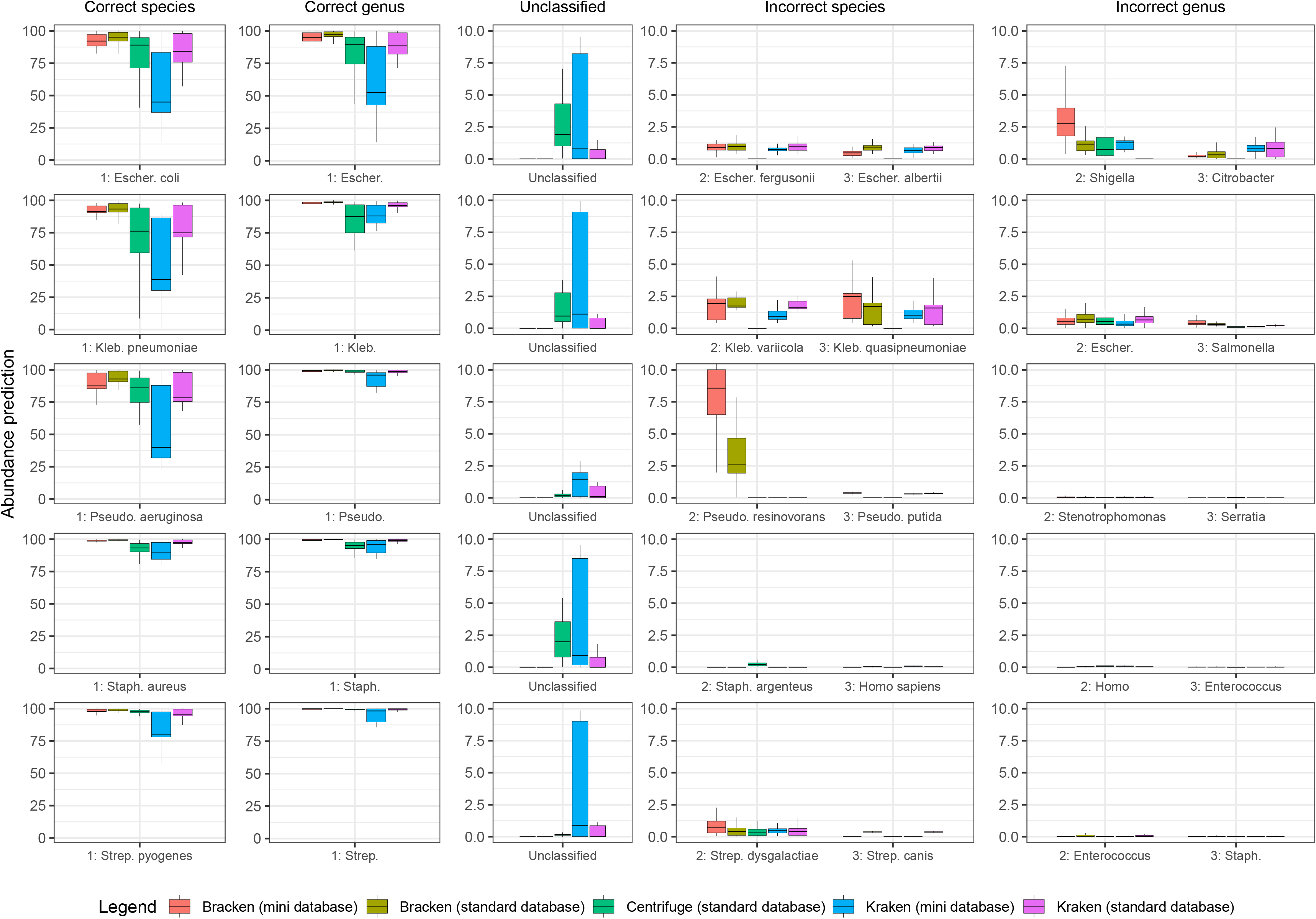
Abundance prediction performance across classifiers for five important blood stream pathogens. The percentage of reads with the correct and incorrect assigned species and genus, and the percentage of reads unclassified are shown.

### Choice of sequencer

Figure 2 details correctly classified, unclassified and misclassified abundances for five common species according to sequencing technology and simulations with and without sequencing errors. Across all 1600 simulated genomes, the theoretical Nanopore error-free performed the best across species, genus, unclassified, misclassified species and misclassified genus abundance [IQR 96.4%:99.5%; IQR 98.5%:100%; IQR 0%:0.1%; IQR 0.1%:0.5% and IQR 0.1%:0.5% respectively] followed by Nanopore including simulated sequencing error [IQR 77.3%:95.7%; IQR 88.8%:98.9%; IQR 0%:3.2%; IQR 0.3%:1.7% and IQR 0%:0.8% respectively], Illumina error-free [IQR 72.8%:97.1%; IQR 93.1%:99.6%; IQR 0%:0.9%; IQR 0.3%:2.1% and IQR 0%:0.7% respectively] and Illumina with sequencing errors [IQR 69.2%:96.9%; IQR 92.1%:99.7%; IQR 0%:1.1%; IQR 0.1%:1.0% and IQR 0.1%:0.3% respectively].

**Figure 2.**
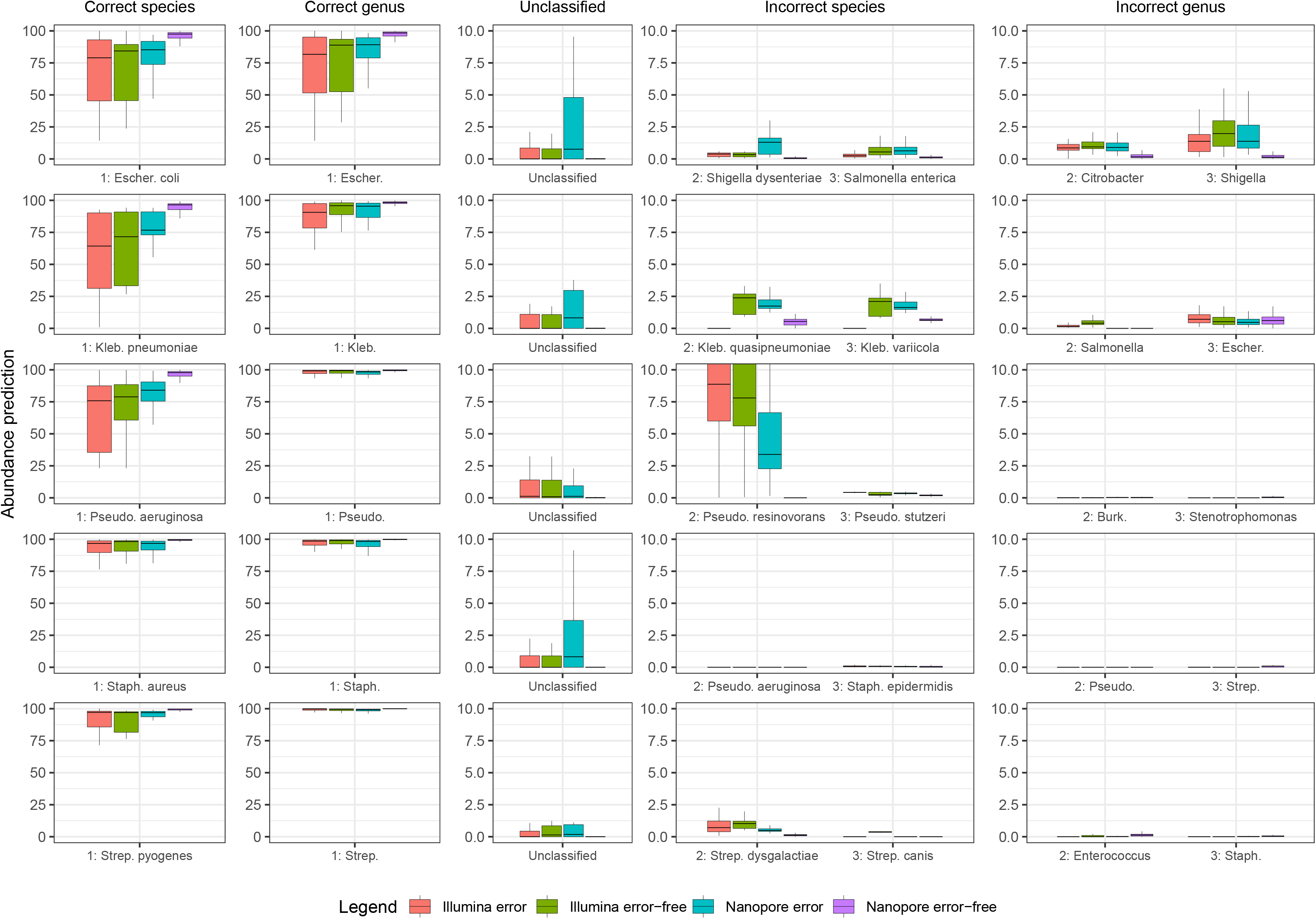
Abundance prediction performance by sequencing device and error profile. The percentage of reads with correct and incorrect classification are shown as well as the percentage of unclassified reads.

### Impact of read length

Figure 3 shows read length vs. proportion of reads correctly classified for both Illumina and Oxford Nanopore sequence simulations, with and without simulated sequencing errors. Performance for *Staphylococcus aureus* and *Escherichia coli* was similar across Illumina error (Range 98.3%-99.2% and 86.7%-94.4% respectively) and error-free profiles (Range 98.4%-99.2% and 86.7%-94.4% respectively), however, differed for Nanopore error (Range 95.6%-99.8% and 83.6%-99.7% respectively) and error-free profiles (Range 98.8%-99.9% and 94.4%-99.9% respectively). Nanopore lengths >13kb performed similarly regardless of error or error-free profile. For *Escherichia coli*, Nanopore error profile read lengths performed similarly to Illumina error-free profile at 3500bp, and performed superiorly at higher read lengths. For *Staphylococcus aureus*, Nanopore error profile read lengths performed similarly to Illumina error-free profile at 10kb, and performed marginally better at higher read lengths.

**Figure 3.**
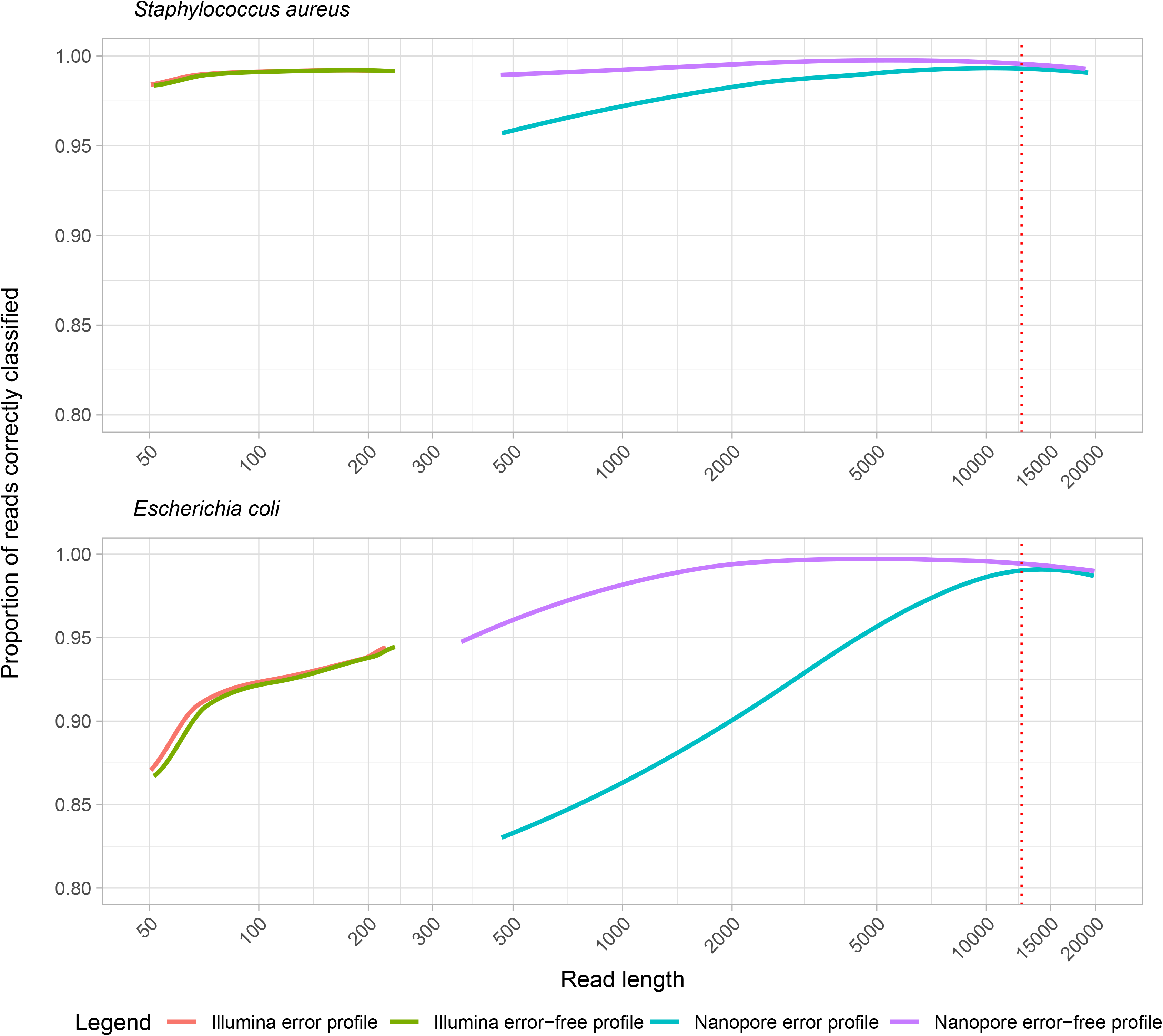
Read length vs. proportion of reads correctly classified.

### Common misclassifications

The top misclassified species for *E. coli* were *Escherichia fergusonii* (median across 100 reference genomes, 0.8%; IQR 0.7%:1.0%; 95^th^-100^th^ percentile 1.8%:20%) and *Escherichia albertii* (median 0.7%; IQR 0.5%:0.9%; 95^th^-100^th^ percentile 1.2%:14.3%), and for genus were *Salmonella* (median 0.5%; IQR 0.3%:0.9%; 95^th^-100^th^ percentile 1.9%:4.8%) and *Citrobacter* (median 0.6%; IQR 0.2%:1%;95^th^-100^th^ percentile 1.7%:21%) respectively. In contrast, *S. aureus* had negligible species and genus misclassification.

Of note misclassified species were more common for *Pseudomonas aeruginosa* which was classified as *Pseudomonas resinovorans* (median 6%; IQR 2.8%:8.9%; 95^th^-100^th^ percentile 11.8%:22.7%), for Streptococcus mitis with misclassification as *Streptococcus pneumoniae* (median 10.7%; IQR 5.9%:16.4%; 95^th^-100^th^ percentile 27.2%:44.7%), and for *Enterobacter cloacae* were *Enterobacter hormaechei* (median 2.1%; IQR 0.8%:25.4%; 95^th^-100^th^ percentile 92%:100%) and *Enterobacter bugandensis* (median 0.1%; IQR 0%:0.4%; 95^th^-100^th^ percentile 92.5%:100%) were identified instead, however, these all belonged to the same species group/complex. Further detailed information on individual species and genus misclassification can be found in Table S3, while Table 3 provides a summary with recommended 95^th^ percentile cut-off value to determine if additional organisms are present, i.e., there is polymicrobial infection.

**Table 3.**
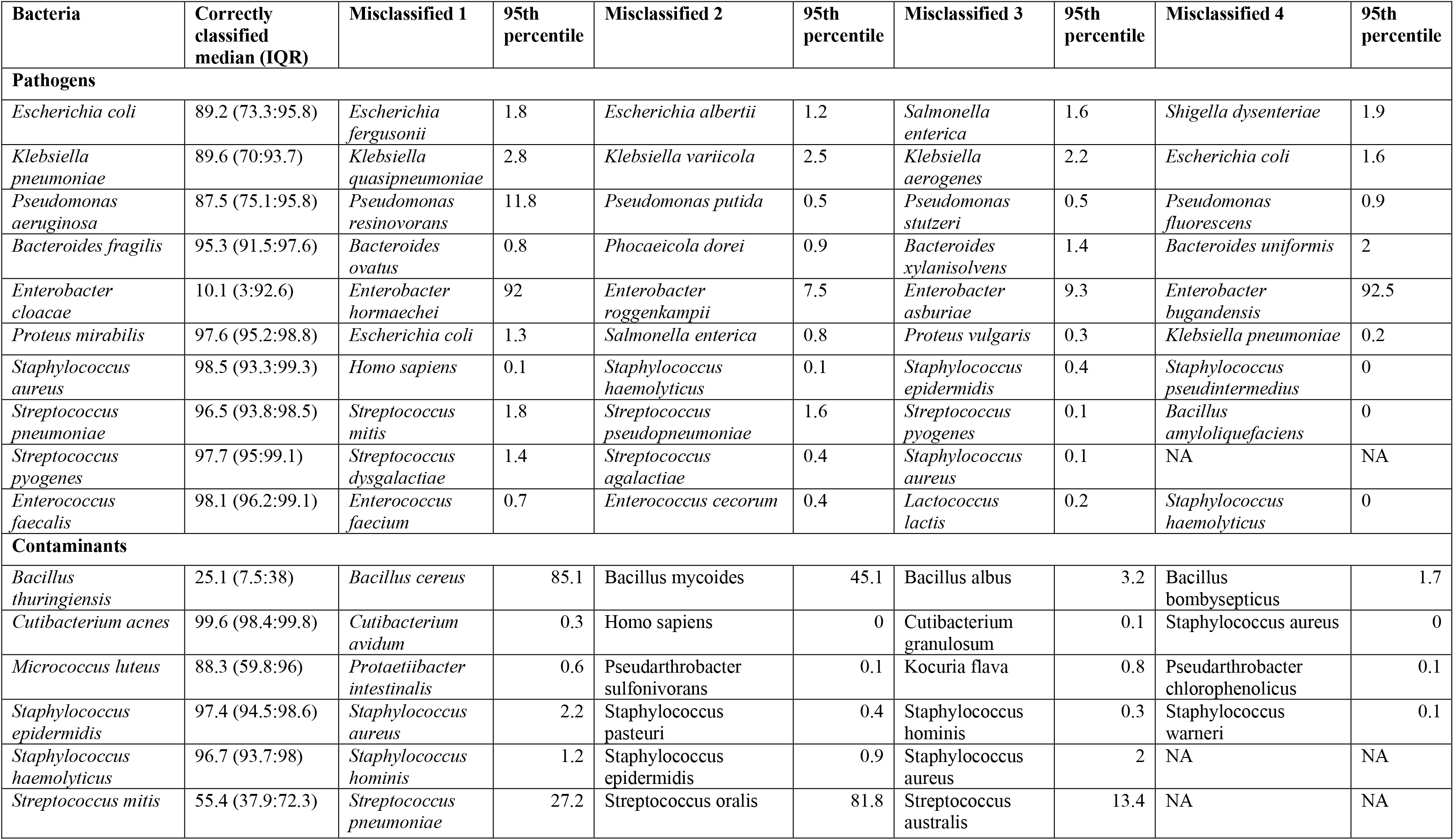
Recommended threshold abundance to call additional true species.

### Relationship between species misclassification and nucleotide identity

Misclassification at the species level was more common when the misclassified species had a higher average nucleotide identity to the true species (Figure 4). However, there were several outliers, where misclassification was greater than expected for the given level of genome-wide nucleotide identity, for example *Bacillus thuringiensis* was misclassified at 25% when *Bacillus cereus* was the true species, and *Streptococcus pneumoniae* at 10% when *Streptococcus mitis* was the true species. This reflects the difficulty distinguishing species within known groups or complexes of bacteria, and potentially reflects that some parts of the genome may have higher nucleotide identity.

**Figure 4.**
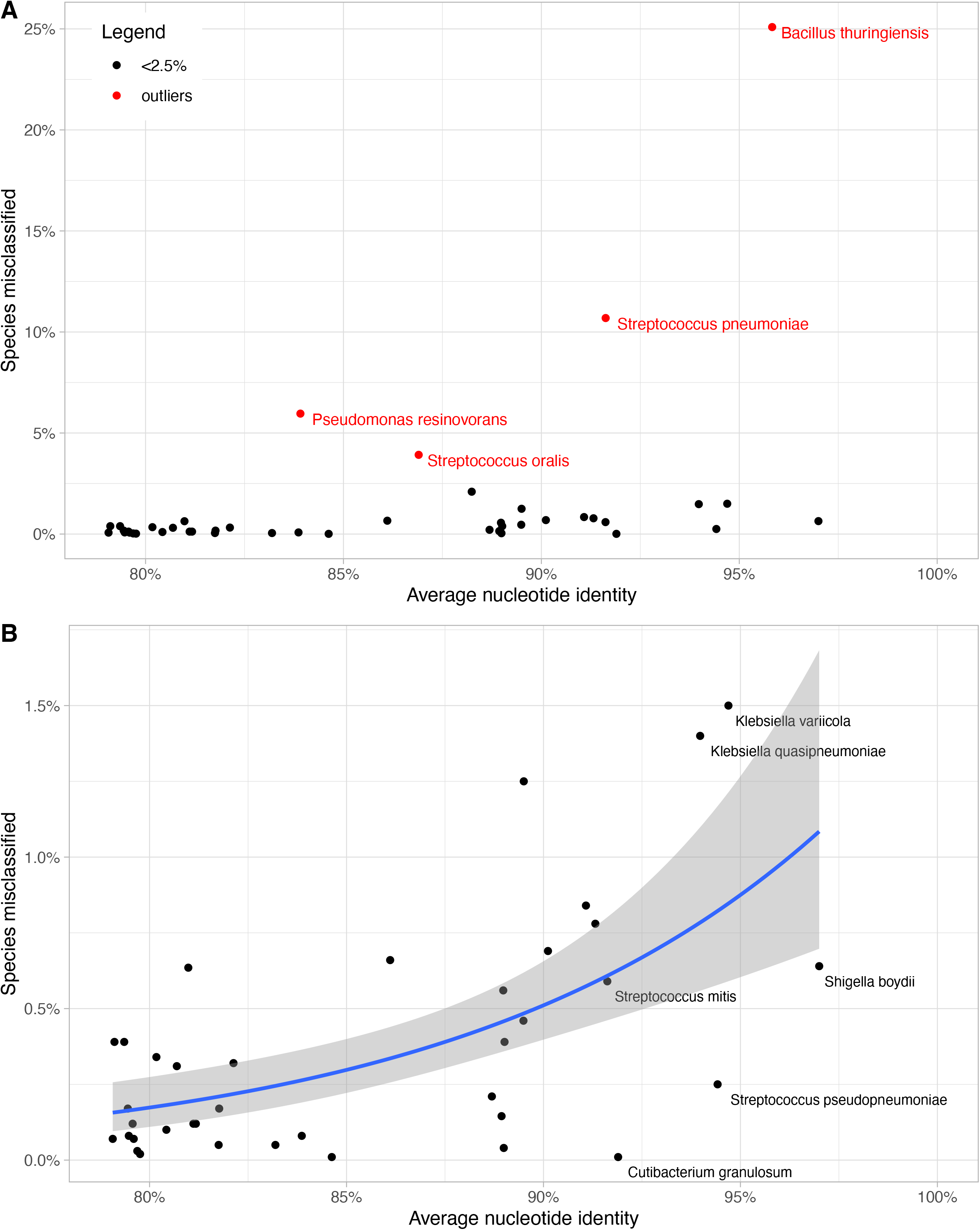
Average nucleotide identity vs. species misclassified. Panel A shows all pairs, and panel B the same plot with the outliers from panel A (shown in red) excluded. For panel B a generalized linear regression line (with a logit link function and binomial family) is plotted (the shaded area indicates the 95% confidence interval). The species corresponding to example outlying points are shown.

### Comparison with real-world data

Illumina simulations correlated with real-world Illumina data (Figure 5). Real-world data from *Escherichia coli, Klebsiella pneumoniae* and *Staphylococcus aureus* performed better than simulated data both when using Kraken2 mini database (percentage of reads classified as the correct species: 45% vs 28.7%; 38.7% vs 19.1%; 90% vs 81.4% respectively) and Kraken2 standard database (84.2% vs 67.1%; 74.9% vs 53.5%; 97.6% vs 96.6% respectively). Similar bioinformatic misclassification existed in both real- and simulated data. Misclassification in *Klebsiella pneumoniae* for example as *Klebsiella quasipneumoniae* was marginally higher in simulated data vs. real-world data in Bracken mini database (2.8% vs 2.5%), Bracken standard database (3.2% vs 1.7%) and Kraken2 standard database (2.2% vs 1.6%).

**Figure 5.**
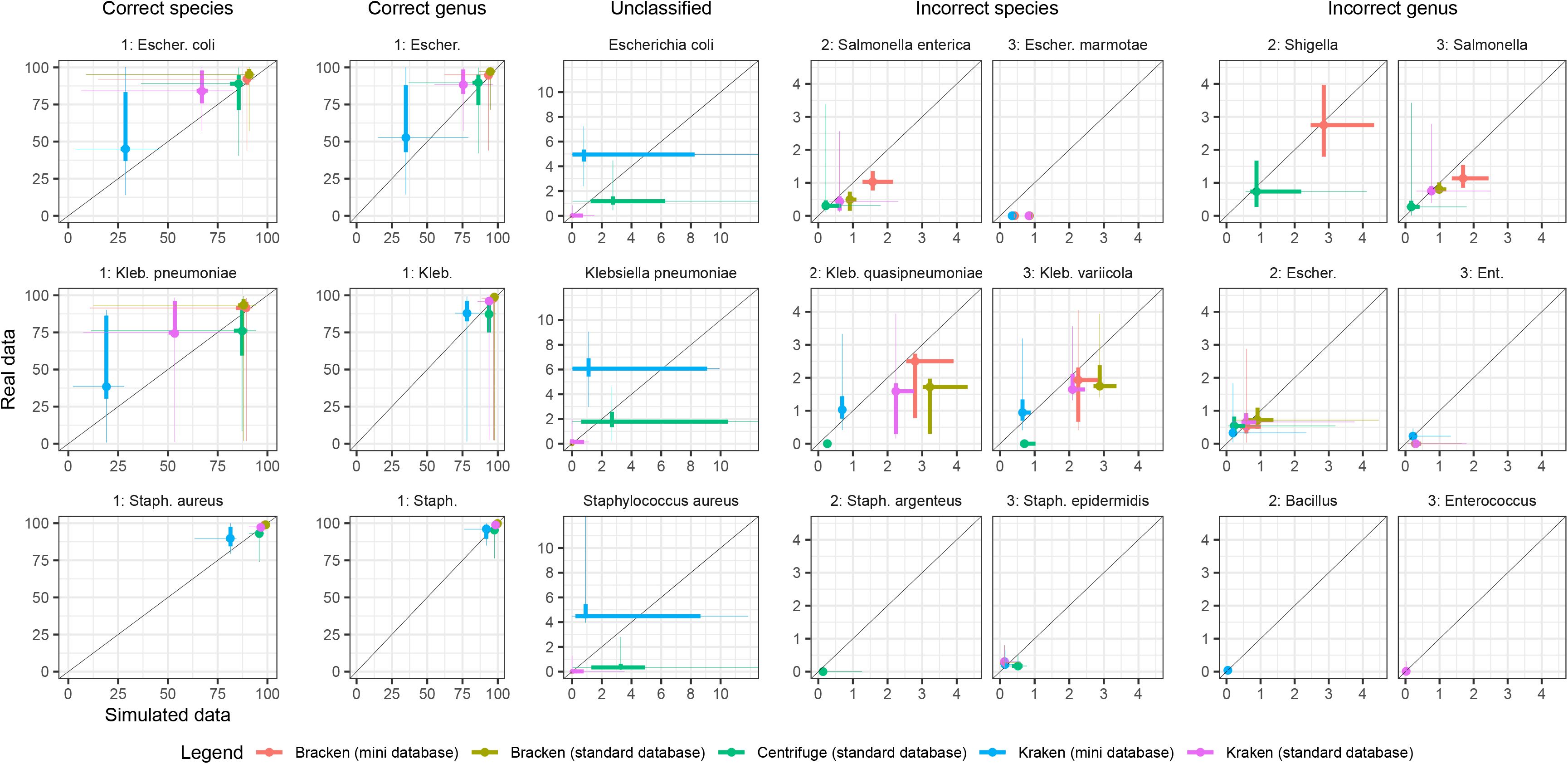
Congruence between Illumina simulated data and real-world data. The diagonal line indicates values with exact correspondence between the simulated a real-world data.

## Discussion

The potential for metagenomic technology to improve diagnostics and patient outcomes have been widely recognized (30), and there is extensive work ongoing to overcome current challenges associated with use of these approaches in a clinical setting (31). In this article, we simulate and use real-world sequencing data to define the intrinsic rates of bioinformatic species misclassification that occur using common sequencing platforms, classification tools and databases. This enables us to quantify the extent of bioinformatic contamination in the classification process to account for this formally in diagnostic workflows, and further guide the selection of optimal bioinformatic tools to maximise classification accuracy.

Overall, across both short and long reads we found that using the Bracken classifier with a “standard” database i.e., larger resulted in the most accurate performance across all species with almost no reads unclassified (Figure 1, and Figure S1). In comparison, Kraken2 with a mini database, a commonly used combination performed worse. Previous studies with Illumina short reads have reported estimates from Kraken2 to be more accurate than Kraken1, KrakenUniq, CLARK and Centrifuge at both the genus and species levels (3). Furthermore, studies have shown with Illumina data that Bracken which re-assigns reads in intermediate taxonomic nodes to the species level or above from Kraken1/2 outputs produce far better abundance estimates (4, 10). Our results confirm that Bracken produces far more accurate abundance estimates for both long- and short-reads, especially when using a standard database. Previous metagenomic studies have rarely used Bracken or standard i.e. larger databases (1), the latter in part because of higher memory requirements, and we encourage future studies to utilise the most accurate tools available where computational resources permit.

To formally account for bioinformatic contamination in clinical diagnostic workflows, we produced the first known catalog of misclassified species from simulating 1,600 genomes representing 10 common pathogens and 6 common blood culture contaminants using different classification tools, databases, and sequencers (Summary catalog Table 3, and detailed catalog Table S3). Additionally, this allows us to specify thresholds for identifying or excluding polymicrobial infections with confidence in metagenomic samples. We used existing real-world Illumina data to show correlation with the Illumina simulator (Figure 5), while the Nanopore simulator used has been benchmarked extensively elsewhere (20). Detailed 95^th^ and 100^th^ percentile cut-off values for other species are available in Table S3 which may be used to determine if there is strong evidence that additional organisms are present, i.e. there is polymicrobial infection or, conversely, if only bioinformatic contamination is present.

Additionally, our results show that misclassification at the species level was more common when the misclassified species had a higher average nucleotide identity to the true species. However, there were several outliers where misclassification was greater than expected for the given level of genome-wide nucleotide identity reflecting the difficulty distinguishing species within known groups or complexes of bacteria. Tools such as DeepMicrobes which use deep learning-based computational framework for taxonomic classification may offer an advantage in these situations where far fewer false positives are produced compared to other tools regardless of different degrees of nucleotide similarity (32).

The advent of long read technologies such as the Oxford Nanopore has significant advantages for metagenomic analysis from the detection of structural variants to improved de novo assembly (33). Furthermore, long-read sequencing of native molecules eliminate amplification bias while preserving base modifications (34). However, these technologies are limited by a higher error-rate affecting 6-12% of all sequenced bases, with a significant fraction of insertion and deletion errors (35). This may affect the success of current classification methods, as there are few algorithms developed to exploit long-read data. However, even after simulating sequencing errors, the longer reads of Nanopore sequencing meant that it still could outperform Illumina sequencing. This finding is supported by previous work (36). Following on from this observation we directly investigated the impact of read length on the proportion of reads correctly classified. Allowing for sequencing error, for *E. coli*, Nanopore reads of 3500bp performed similarly to Illumina reads and performed better at higher read lengths. For *S. aureus* Nanopore with 10kb reads performed similarly to Illumina, and performed marginally better at higher read lengths. Our findings have several implications: As technology and accuracy for long-read sequencing advances, we expect classification accuracy to significantly improve. Furthermore, filtering shorter reads (at least <3500bp) from error-prone Nanopore sequencing can provide performance either equivalent or superior to standard illumina sequencing.

In conclusion, our findings highlight the pertinent issue of the presence of bioinformatic contamination and how this varies by the combination of sequencer, classifier and database used. While we have recommended Bracken using a standard database to be utilized in metagenomic workflows, we have produced a misclassification catalog whereby misclassification can be accounted for using any combination of sequencers, classifiers or databases benchmarked in this study. Benchmarking metrics should inform the choice and implementation of metagenomic sequencing tools for both research and clinical applications.

## Acknowledgments

Dr. Sam Lipworth for providing real-world sequencing data to be used in this analysis

## Supplementary figures

**Figure S1. Abundance prediction performance across classifiers for five common blood stream contaminants**. The percentage of reads with the correct and incorrect assigned species and genus, and the percentage of reads unclassified are shown.

**Figure S2. Real-world abundance classification performance of pure isolates from Illumina sequencing**.

## Supplementary tables

**Table S1. Logistic regression model for performance of genus abundance classification by sequencer, classifier, database and bacterial species.**

**Table S2. Predicted abundance classification for all included species.**

**Table S3. Detailed threshold abundance recommended to call additional true species and genus.**

